# Elevated levels of active Transforming Growth Factor β1 in the subchondral bone relate spatially to cartilage loss and impaired bone quality in human knee osteoarthritis

**DOI:** 10.1101/2021.09.13.459432

**Authors:** Dzenita Muratovic, David M. Findlay, Ryan D. Quarrington, Xu Cao, Lucian B. Solomon, Gerald J. Atkins, Julia S. Kuliwaba

## Abstract

**Objective:** Over-activity of transforming growth factor β1 (TGFβ1) in subchondral bone has a direct causal role in rodent models of knee osteoarthritis (OA), which can be blocked by TGFβ1 neutralisation. In this study, we investigated whether the spatially distributed level of active TGFβ1 in human subchondral bone associates with the characteristic structural, cellular and molecular parameters of human knee OA.

**Design:** Subchondral bone samples (35 OA arthroplasty patients, aged 69±9 years) were obtained from regions below either macroscopically present or denuded cartilage. Bone samples were processed to determine the concentration of active TGFβ1 (ELISA) and gene-specific mRNA expression (RT-PCR). Synchrotron micro-CT imaging was utilised to assess the bone microstructure, bone mineralization, the osteocyte lacunar network and bone matrix vascularity. Finally, samples were histologically examined for cartilage OARSI grading, quantification of tartrate resistant acid phosphatase positive cells and bone marrow micro-vasculature.

**Results:** Subchondral bone below severely degenerated/depleted cartilage, characterised by impaired bone matrix quality due to sclerotic microarchitecture, disorganised collagen, high heterogeneity of the mineral distribution, contained increased concentrations of active TGFβ1, compared to adjacent areas with more intact cartilage. In addition, increased levels of active TGFβ1 related directly to increased bone volume while increased OARSI grade associated directly with morphometric characteristics (size, shape and orientation) of osteocyte lacunae.

**Conclusion:** These results indicate that increased active TGFβ1 associates spatially with impaired bone quality and the disease severity of human OA. This study therefore suggests that TGFβ1 could be a therapeutic target to prevent or reduce human disease progression.

## Introduction

Osteoarthritis (OA) is a painful degenerative disease of articulating joints and is the most common single cause of pain and disability after middle age for both men and women [1]. A lack of understanding of the underlying mechanisms of OA has prevented the development of disease modifying treatments.

While OA is characterised by progressive degenerative damage to articular cartilage, it is now widely accepted that the disease affects the whole joint. Characteristic changes that occur in the sub-chondral bone (SCB) in OA depend on the stage of disease progression [2] and are not uniform across the joint [3]. Seen most prominently beneath areas of cartilage degeneration and loss, there is increased bone remodelling, bone sclerosis, and vascular changes [2, 4-6].

Transforming Growth Factor β1 (TGFβ1) has multiple essential roles in development, homeostasis, and repair in both cartilage and SCB. In bone, TGFβ1 acts as a chemoattractant to recruit osteoprogenitor cells to sites of new bone formation, where it stimulates their proliferation and matrix production [7]. TGFβ1 also acts directly on osteoclast precursors to attract them to sites of bone resorption, and on osteoblasts to increase their expression of osteoclast regulatory molecules, including RANKL and M-CSF [8]. Osteoclasts in turn release and activate latent TGFβ1 stored in the bone matrix [9], which recruits osteoblasts, coupling bone resorption with bone formation [10]. Moreover, TGFβ1 acts on osteocytes to regulate bone volume indirectly [11] and bone matrix material properties directly [12]. TGFβ1 also acts on endothelial cells, having critical roles in both normal and pathological blood vessel formation [13]. These important actions of TGFβ1 in bone show the potential for its involvement in skeletal disease, and inappropriately high levels of active TGFβ1 have been linked to osteogenesis imperfecta [14], tumour-induced osteolytic bone disease [15], and OA [16].

OA studies in mice provide evidence that increased levels of active TGFβ1 in cartilage and SCB compromise functional integrity of those tissues and lead to pathological changes that are causative of OA [17]. Mice with transgenic expression of active TGF-β1 showed spontaneous OA-like changes in the SCB and degradation of the overlying cartilage, strongly suggestive of cross-talk between these tissue compartments [18]. Hallmarks of increased TGFβ1 activity in the SCB were: (i) increased numbers of nestin-positive mesenchymal stem cells and osterix-positive cells (osteoprogenitors) in the bone marrow; (ii) increased angiogenesis; (iii) the appearance of osteoid islands; and, (iv) lesions imaged by magnetic resonance imaging (MRI) similar to bone marrow lesions (BMLs) seen in human knee OA. Importantly, the early structural changes in subchondral bone alter the distribution of stress on the articular cartilage and directly result in its degeneration. Neutralising TGFβ1, either locally in the SCB or systemically, attenuated OA changes in the SCB and articular cartilage in mouse and rat OA models, suggesting that TGFβ1 acting in the SCB is causal of OA. [16-18].

The accumulated evidence indicates an urgent requirement to investigate whether TGFβ1 might play a similar role in the human disease. Such studies are an essential precursor to testing whether TGFβ1 neutralisation might be effective clinically to prevent or slow knee OA disease progression. Thus, the aim of this study was to investigate whether the level of active TGFβ1 protein in the subchondral bone associates with the structural, cellular and molecular parameters that are characteristic of human knee OA.

## Materials and Methods

### Clinical Specimens

Tibial plateaus were collected from 35 patients (15 males aged 66±9 years and 20 females aged 70±8 years), undergoing total knee replacement surgery. Inclusion criteria were a diagnosis of end-stage knee OA based on radiographic imaging, and symptomatic disabilities. Exclusion criteria were knee OA due to previous trauma, rheumatoid arthritis, osteoporosis, malignancy, metabolic bone disease, and medication that may have affected bone metabolism, such as steroids. Written consent was obtained for all subjects and the study received prior approval from the Human Research Ethics Committee at the Royal Adelaide Hospital and The University of Adelaide, South Australia.

Since changes in subchondral bone in human OA are not uniform across the whole tibial plateau, several trabecular bone samples were collected from regions with visually present and relatively intact cartilage (CA+) and adjacent regions with severely degenerated/depleted cartilage (CA-), (Fig. 1A). Specifically, two bone core biopsies (8×8×7 mm) from CA+ and CA- regions were sampled immediately after surgery and were cleaned from bone marrow by several wash in sterile PBS to determine concentration of TGFβ1 in subchondral bone and to quantify mRNA levels. Detailed protocols are described in the Supplementary Methods.

**Figure 1.**
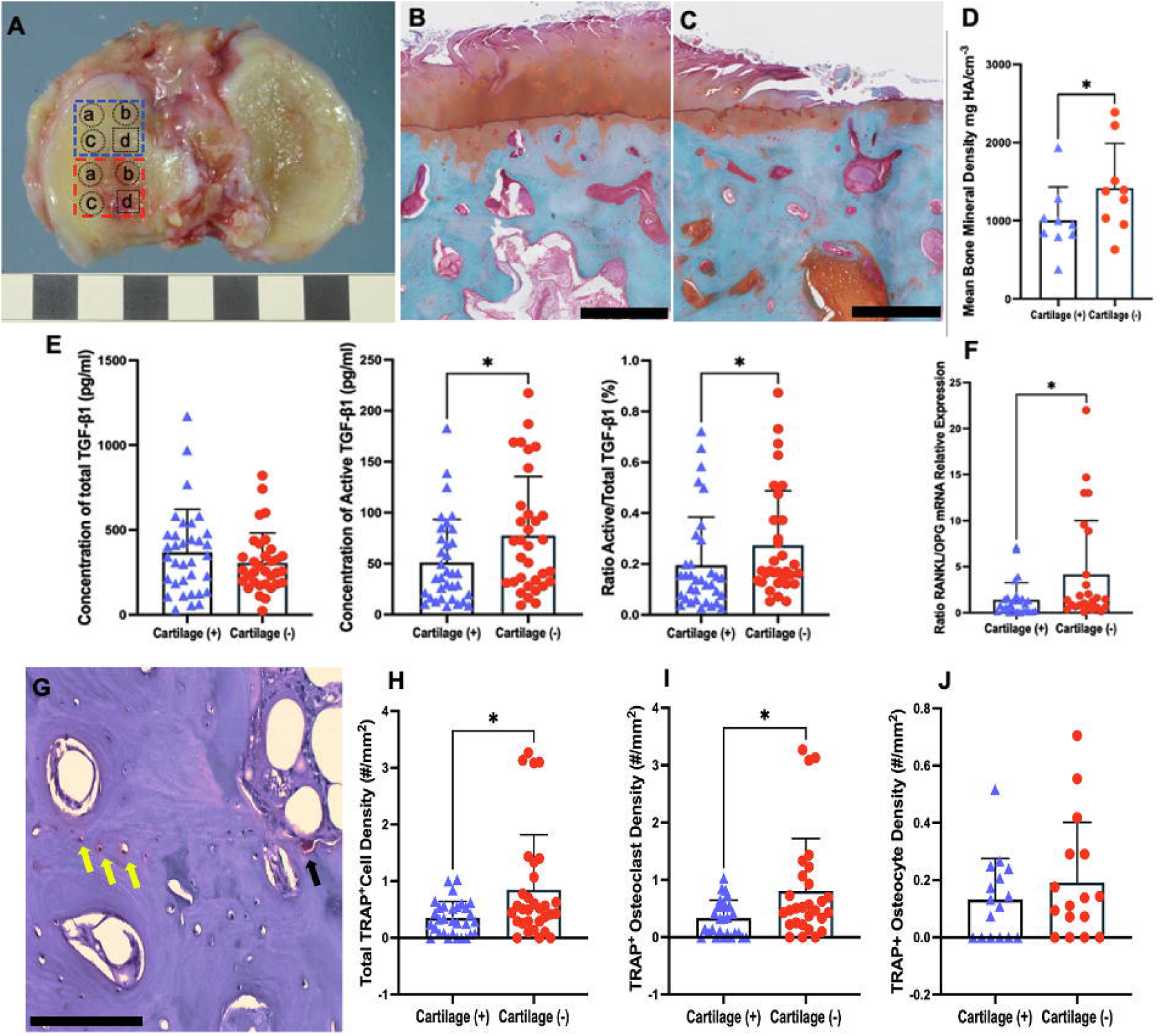
Gross view of an osteoarthritic tibial plateau (female, age 73 years); blue square shape showing region with macroscopically intact cartilage; red square shape showing region with severely degraded cartilage. a) Bone core used to determine the concentration of active and total TGFβ1 by protein assay; b) Bone core used to quantitate relative expression of mRNA by real-time PCR; c) bone core for a subset of 9 subjects scanned by synchrotron radiation micro-computer tomography d) tissue used for histological examination. Representative Safranin O Fast Green stained sections in Cartilage+ (B) and Cartilage-regions (C) to determine mean histological OARSI grades (D). Concentration of total TGF β1, active TGFβ1 and the ratio of active/total TGFβ1 in SCB in CA+ and CA- regions, respectively (E). The ratio of RANKL/OPG mRNA relative expression (F). Histochemical TRAP staining, yellow arrows indicate TRAP+ osteocyte, black arrow indicates TRAP+ osteoclast (G). Total TRAP+ cell density (TRAP+ osteoclast + TRAP+ osteocytes), TRAP+ osteoclast cells TRAP+ osteocytes density (H). Each point represents an individual sample. Data shown as median (25th, 75th percentiles). * p < 0.05.

### Histomorphometry

Blocks of osteochondral tissue (10×10×8 mm) directly adjacent to the cores sampled for molecular measures, were cut from both regions using a low-speed diamond wheel saw (Model 660, South Bay Technology, San Clemente, CA, USA), and subjected to histological examination; Safranin O Fast Green was used to evaluate the degree of degenerative changes in osteochondral unit (cartilage and bone) [19]; Haematoxylin and Eosin (H&E) was used to quantify osteocyte density in the trabecular bone and arteriolar density in the subchondral bone marrow. Ploton silver staining was used for visualisation of the osteocyte lacunocanalicular network [12, 20]. Histochemical TRAP staining was used to quantify TRAP+ cells within the subchondral bone [21].

Polarised-light microscopy was performed on sections stained by picrosirius red stain [12, 22]. In order to estimate the local orientation of the collagen fibres, we used OrientationJ, an ImageJ plug-in, as described previously [23].

### Synchrotron radiation micro-computer tomography (SRμCT)

An additional bone core biopsies from CA+ and CA- were sampled from a subset of 9 patient, for imaging by synchrotron micro-CT. To ensure that evaluation of bone quality by synchrotron micro-CT corresponds to overall group values, we selected core biopsies from 4 females (aged 74.5 ±6 years) and 5 males (aged 66±6) with OARSI grade and mean concentration of total and active TGFβ1 corresponding to means of the overall group.

Synchrotron tomographic X-ray experiments were carried out at the X02DA TOMCAT beamline of the Swiss Light Source (SLS) facility at the Paul Scherrer Institute (PSI). Detailed scanning protocol is described in supplementary method. Using CT-An analyser (SkyScan, Kontic, Belgium), cylindrical regions of interest (size 3×3×1 mm) with approximate volume of 7mm^3^, containing the trabecular bone from CA+ and CA- regions was used for analysis of trabecular bone microstructure, degree of mineralization, density and morphometry of osteocyte lacunae and density of vascular canals in bone matrix.

3D morphometric trabecular bone microstructure parameters included: bone volume fraction (BV/TV), trabecular thickness (Tb.Th), trabecular number (Tb.N), and trabecular separation (Tb.Sp).

The degree of the bone mineral density was evaluated based on the original reconstructed images using custom written MATLAB code. The grey level corresponding to the bone mineral peak was extracted. As previously published, the x-ray attenuation coefficient (grayscale values) for each voxel was divided by the mass attenuation coefficient of bone (4.001 cm2/g at 20 keV) to directly calculate bone mineral density distribution and the peak calcium content. To quantitatively determine the distribution of the mineral content, mean calcium content (Ca mean), heterogeneity of the calcium distribution (Ca width), most frequent calcium content (Ca peak), amount of poorly mineralized bone (Ca low), and amount of highly mineralized bone (Ca high) as previously explained [24].

Osteocyte lacunae were distinguished in bone images using 2 step Otsu thresholding. Based on a previously published report, a despeckling (denoising) function was used to remove structures less then 30 µm^3^ (believed to be noise) and above 2000 µm^3^ (believed to be vascular canals) and the remaining closed pore like structures were defined as osteocyte lacunae [25]. Osteocyte lacunar parameters; *lacunar density (La*.*Dn, #/mm*^*2*^*), lacunar volume (La*.*V, µm*^*3*^*) orientation of* lacunae (°), *Sphericity (Sph*) [26], and Structure model index (SMI) [27] were obtained using individual object analysis.

The vascular canal network within bone matrix at the micro-level was determined by the following parameters: canal density (canal number/bone volume, #/mm^3^), total canal volume (*µ*m^3^) and mean vascular canal diameter(*µ*m) [28].

### Statistical analysis

The Shapiro-Wilk test was performed to assess the distribution of data. Comparisons between CA+ region and CA- regions were evaluated using paired, two-tailed Student’s t-test for parametric data or Wilcoxon test for non-parametric data. These analyses were performed using GraphPad Prism (GraphPad Software, Inc., San Diego, CA, USA). Parametric data are presented as mean ± standard deviation, and non-parametric data as median ± interquartile range. Statistical significance was set at α = 0.05. Further, linear mixed-effects models were used to investigate a direct association between levels of concentration of active TGFβ1, OARSI score, and parameters describing structural, cellular, and molecular tissue characteristics in subchondral bone, adjusting for sex, age, and cartilage quality in the analysed region. These analyses were performed using SPSS v26 (IBM, Chicago, IL, USA).

## Results

### Degree of osteochondral degeneration relates spatially to concentration of active TGFβ1 in human subchondral bone

OARSI grades ranged between 2 and 4.5 (median=3.5) in CA+ regions, and between 3 and 6.5 (median=6) in CA- regions, indicating that regions labelled as CA- corresponded to severe degeneration of osteochondral tissue, while CA+ regions corresponded to moderate degeneration (*p*<0.001) (Fig. 1B-D). Quantitative measurement of TGFβ1 protein in subchondral bone indicated no significant difference in total TGFβ1 concentration between CA- and CA+ regions. However, significantly higher mean concentrations of active TGFβ1 (p=0.02) and mean ratio of active/total TGFβ1 (p=0.01) were found in trabecular bone below severely degenerated cartilage when compared to bone below moderately degenerated cartilage (Fig. 1E).

Evaluation of the expression of genes associated with bone resorption (*TRAP/ACP5, RANKL/TNFSF11* and *OPG/TNFRSF11B*) and bone formation (*DMP1, TNAP, OCN/BGLAP, COLA1*), as well as *TGFB1* and *SMAD3*, indicated significantly higher mean expression of the *RANKL*/*OPG* mRNA ratio (*p* = 0.03) in CA- regions compared to CA+ regions (Fig. 1F). These results are consistent with previously described uncoupled bone resorption in human hip OA [29]. However, no significant differences were found between the two regions in the relative gene expression of the other genes examined (*Supplementary Fig.1*).

TGFβ1 is produced and stored in bone matrix as a latent complex during bone formation and released as an active ligand in response to mechanical stimuli during osteoclastic bone resorption. Quantitative analysis of TRAP^+^ osteoclasts, indicated significantly more TRAP^+^ osteoclasts (p=0.03), in CA- regions compared to CA+ regions. Interestingly, TRAP staining revealed that in addition to TRAP^+^ osteoclasts, TRAP^+^ osteocytes were also present in the bone tissue of both CA+ and CA- regions (Fig. 1G). TRAP^+^ osteocyte density was not significantly different between the two regions, (Fig. 1H-J).

### Bone matrix quality is distinctively different between CA- and CA+ regions and relates spatially to zones with increased levels of active TGFβ1

Animal studies have demonstrated that TGFβ1 signalling plays a direct role in the regulation of bone remodelling in response to mechanical demands, and consequently affects properties and quality of bone tissue [30]. As expected, subchondral trabecular bone from the CA- regions was significantly more sclerotic, with 38% increase in bone volume (*p*<0.05) and 50% higher trabecular number (*p*<0.05), along with 11% decreased trabecular separation (*p*<0.05) compared to CA+ regions (Fig. 2A-C).

**Figure 2.**
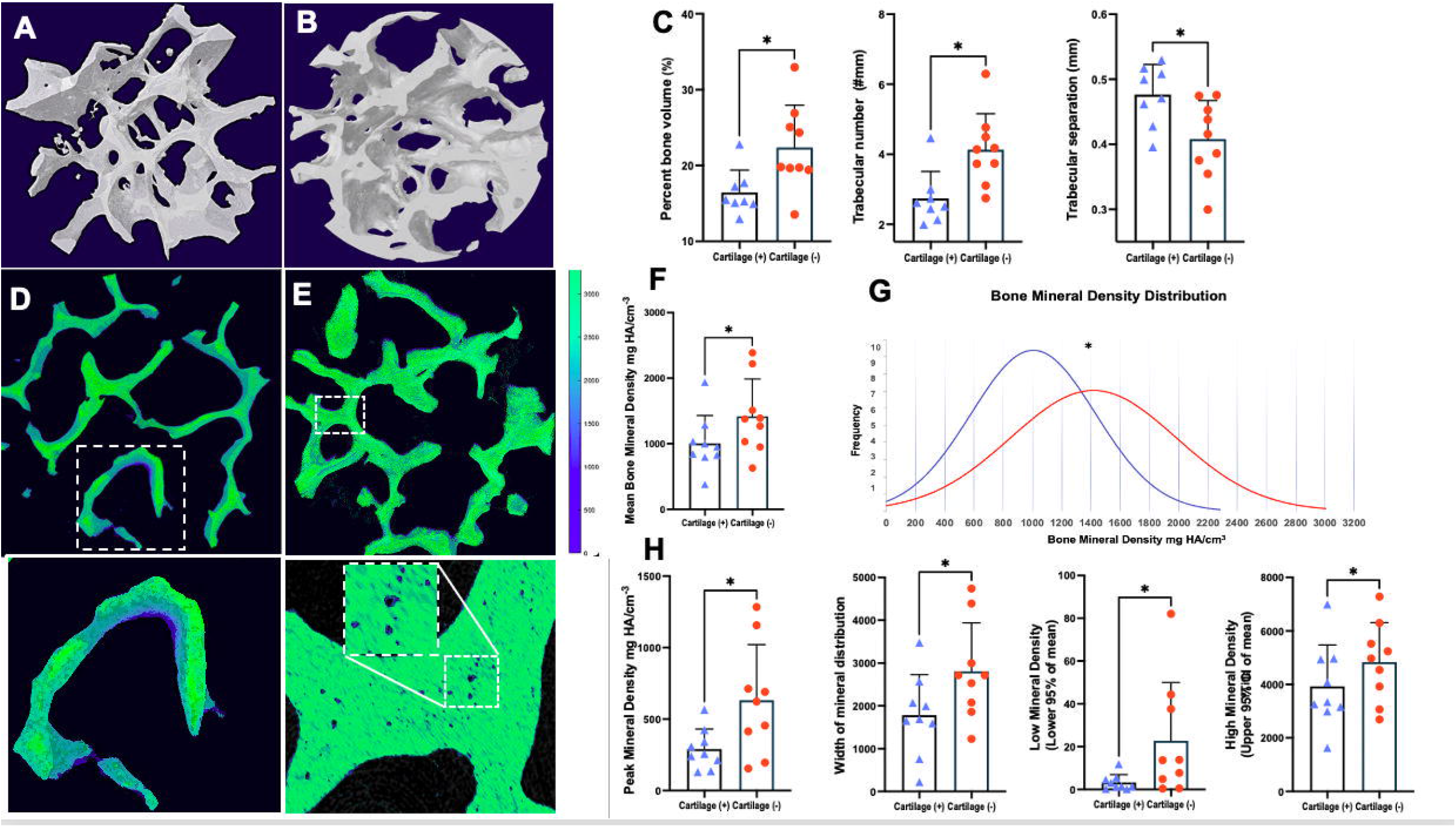
Reconstructed 3D models from Synchrotron Radiation X-ray micro- computed tomography images of subchondral bone microstructure in (A) Cartilage+ and (B) Cartilage-regions. Subchondral bone below severely degraded cartilage is characterised by significantly increased bone volume, trabecular number and decreased trabecular separation (C), Data are presented as mean ± SD, *p<0.05. 3D reconstructed colour coded images of mineral density distribution in CA+ and CA- regions (D&E). Mean bone mineral, mineral content as a frequency distribution (histogram) of grey levels, most frequent mineral content (Ca peak), heterogeneity of the mineral distribution (Ca width), amount of poorly mineralized bone (Ca low) and amount of highly mineralized bone (Ca high) between CA+ and CA- regions (F-H). Data are presented as mean ± SD, Scalebar 100 µm, *p<0.05.

A significant increase in mean bone mineral density was found in CA- regions (*p*=0.02) relative to CA+ regions, (Fig. 2 D-F). Additionally, bone matrix in CA- regions was characterised by increased peak mineral density (*p*=0.01), and increased heterogeneity of the mineral distribution (mineral density distribution width), (*p*=0.03). Specifically, we found that bone in CA- compared to CA+ regions contain significantly greater areas of poorly mineralized bone (low mineral density, *p*=0.01) but also significantly increased amounts of highly mineralized bone (high mineral density, *p*=0.03), (Fig 2 G&H).

3D reconstructed colour coded images of mineral density distribution show strikingly different patterns of mineralisation between CA+ and CA- regions, (Fig. 2D&E). Ca+ regions displayed a more organised pattern, perhaps due to lower bone formation rate and thus more homogenous mineralisation across the bone tissue. CA- regions contained numerous less mineralised bone packets distributed diffusely throughout the bone possibly due to dynamic and less organised bone formation. We also noticed that in CA- regions, the osteocyte perilacunar bone matrix was characterised by low mineralisation (Fig. 2E), most likely reflecting an ongoing active perilacunar remodelling in the CA- bone. Thus, specific mineralisation density distribution in CA- regions likely indicate increased and uncoupled bone remodelling, associated with osteocyte perilacunar remodelling.

Consistent with the mineral distribution, polarised-light microscopy confirmed that CA- regions were composed of more randomly arranged collagen fibrils, compatible with newly deposited, woven bone matrix, while in CA+ regions collagen fibrils were more organised and with preferential orientation (*p*=0.03), (Fig. 3A-E). Taken together, the data describing bone matrix characteristics indicated that the integrity of bone tissue in CA- regions was significantly impaired, and these changes associated spatially with increased levels of TGFβ1.

**Figure 3.**
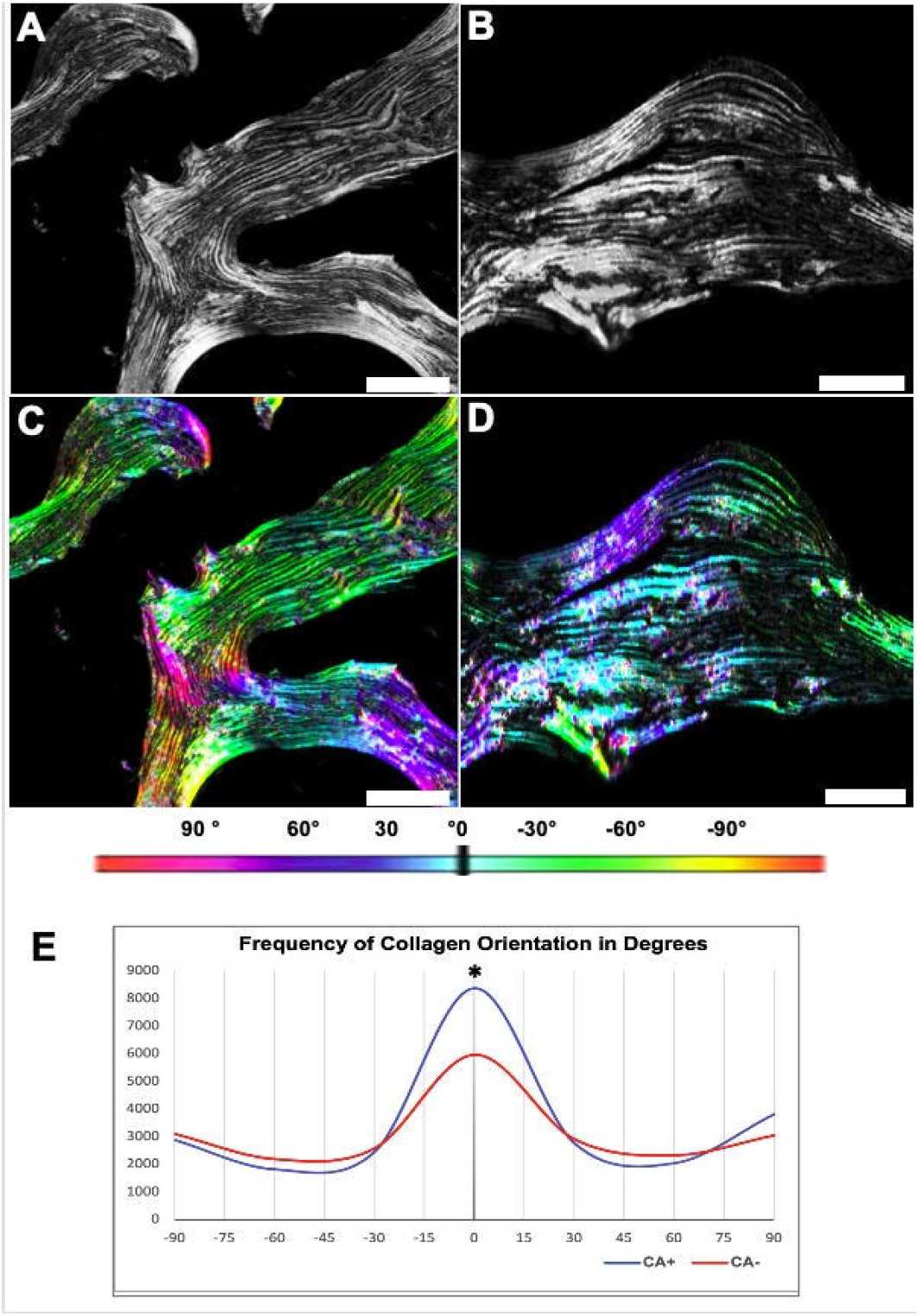
Binarized images of sections stained with Picrosirius Red stain at 40x magnification (A&B) and color-coded orientation of fibres (C&D) according to the colorimetric scale (−90°-90°). E) Collagen fibre organisation in subchondral bone regions below severely degraded cartilage (CA-) was less organised, compared to regions below moderately degraded cartilage (CA+). Scalebar 100 µm, *p<0.05.

### Osteocyte-lacunocanalicular network

SR-µCT analysis of osteocyte lacunae (Fig. 4 A-C) indicated significantly higher osteocyte lacunar density (p=0.02) in bone below CA- regions. In addition, osteocyte lacunae in CA- regions were characterised by increased lacunar volume (p=0.02) and altered shape that was less elongated (SMI; p=0.04), and less spherical (Sph; p=0.02) compared to lacunae in CA+ regions, (Fig. 4D). The orientation of lacunae in CA- regions was also significantly different compared to the orientation of lacunae in CA+ regions (p=0.04), (Fig. 4D). These differences in shape and orientation of the osteocyte lacune between the regions are possibly indicators of the transition process from woven to lamellar bone [24]. In addition, these results are also suggestive of dynamic active perilacunar remodelling in CA- regions, which has been reported to be a direct consequence of local osteocyte-derived TGFβ signalling [7, 12, 31-33]. Further, quantitative histological assessment (Fig. 4E-G) also indicated increased osteocyte density (21%) in the CA- regions relative to CA+ (p=0.0009), suggesting that CA- bone had been more recently remodelled [24]. Consistent with this, the ratio of empty over total lacunae were significantly higher in CA+, compared to CA- (p=0.0004). Using the Ploton silver stain (Fig. 4H&I), no quantitative differences were found for canalicular number and length of canaliculi between the CA+ and CA- regions.

**Figure 4.**
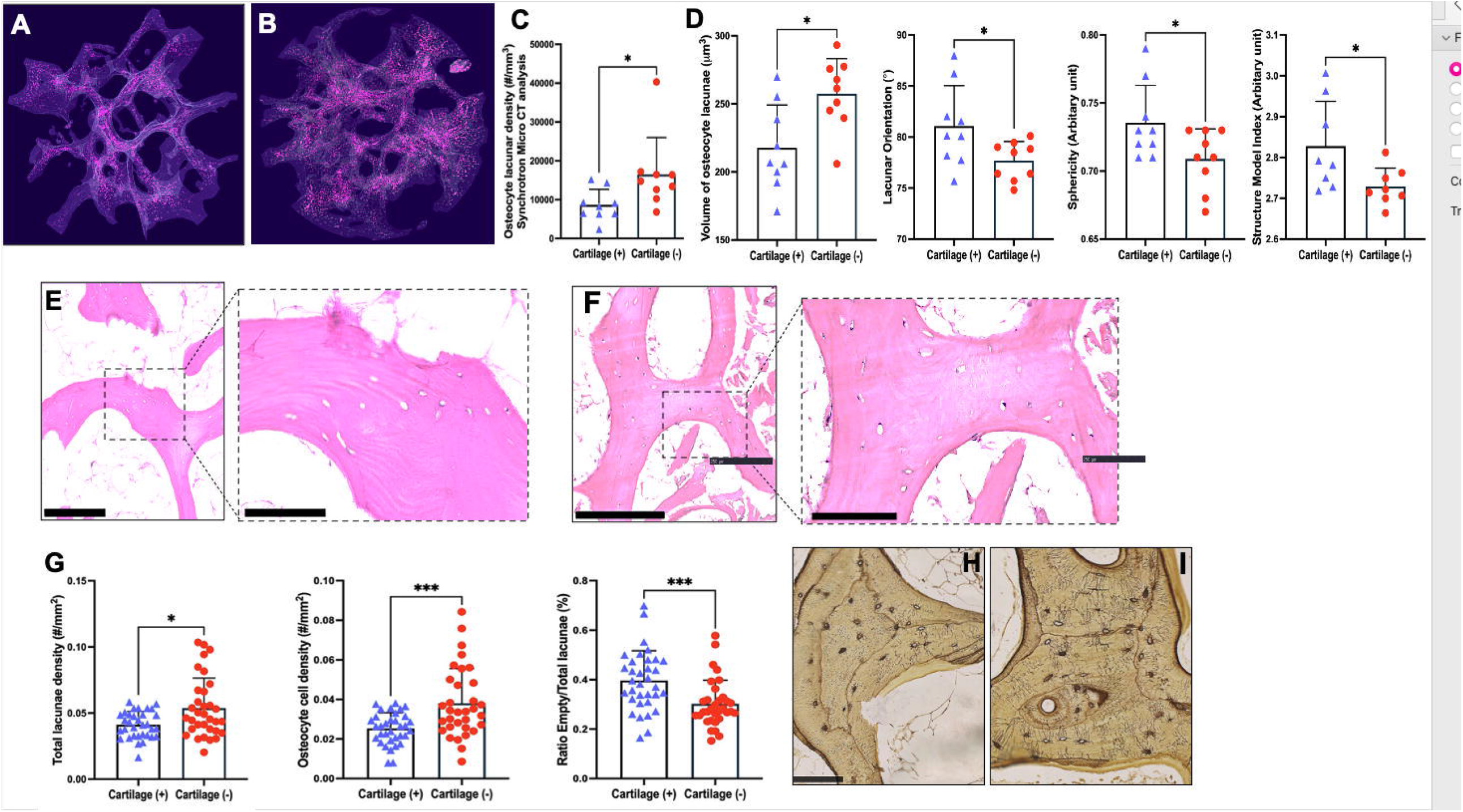
A&B, Synchrotron Radiation X-ray micro-computed tomography images of subchondral bone reconstructed in 3D (light grey) and osteocyte lacunae (pink) in CA+ and CA- regions. Haematoxylin and Eosin was used to quantify osteocyte density in CA+ and CA- (E&F) regions. Ploton silver stain was used for visualisation of the osteocyte lacunocanalicular network CA+ and CA- (H&I). Data shown as median (25th, 75th percentiles), *p<0.05, ***p<0.0005.

Analysis of vascular density indicated significantly increased vascularity in both bone matrix, showing increased vascular channel density (Fig. 5A-C, p=0.02), and bone marrow showing increased arteriolar density (Fig. 5D-F, p=0.002) in CA- compared to CA+ regions.

**Figure 5.**
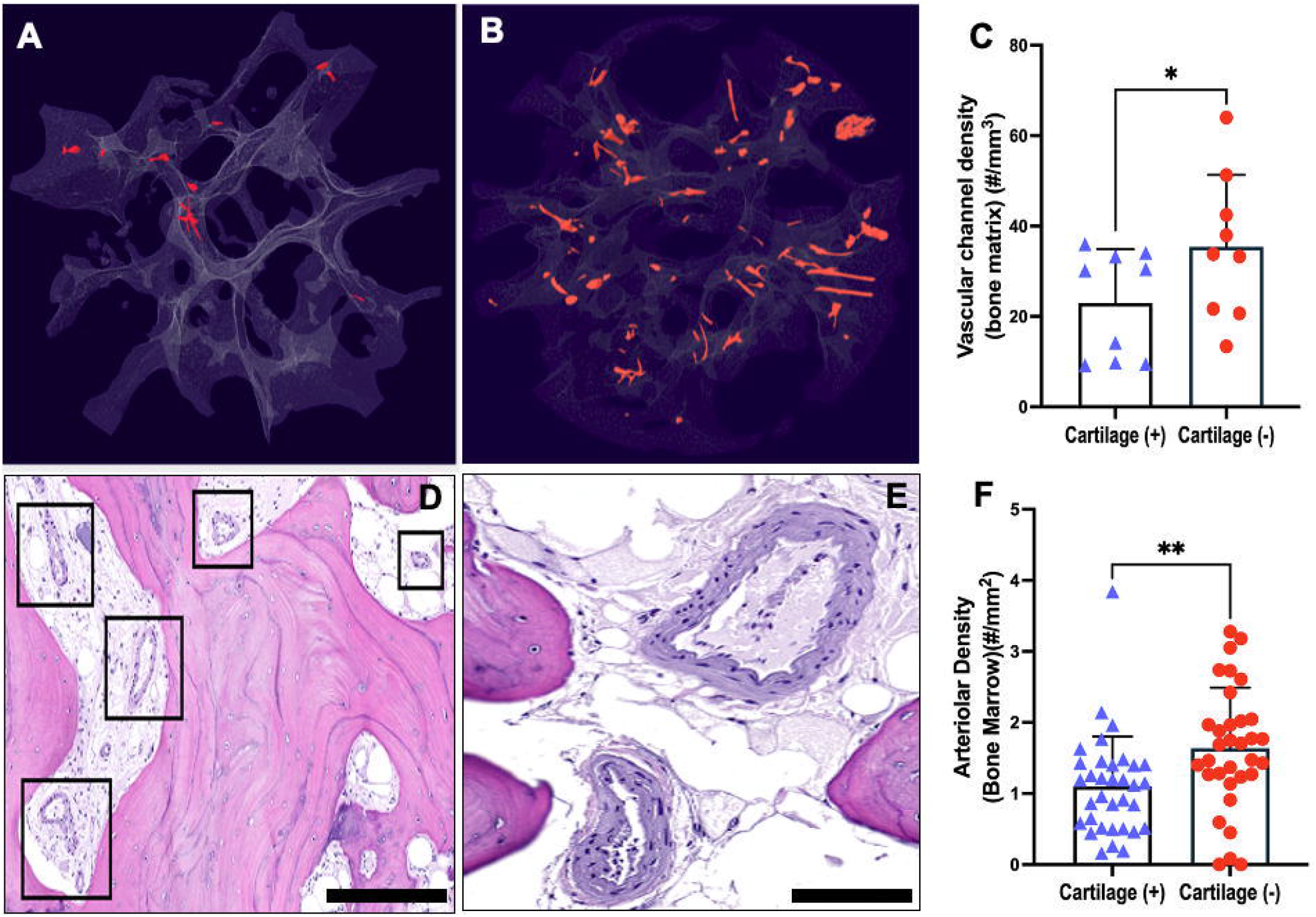
A Synchrotron Radiation X-ray micro-computed tomography images of subchondral bone reconstructed in 3D model (light grey) and 3D model of Vascular channels(red) within bone matrix in Cartilage+ and Cartilage- (A&B) regions. Quantitative analysis showed significantly increased density of vascular canals within bone matrix (C). Thick wall arterioles in bone marrow (D&E) between the subchondral trabeculae, indicated by a black square (D). and Quantitative analysis indicating increased arteriolar density in bone marrow in CA- compared to CA+ regions (F).

### Association between concentration of active TGFβ1 and OARSI score and parameters describing structural, cellular, and molecular tissue characteristics, adjusting for sex, age and bone region

Linear mixed-effects models indicated a direct relationship between the increased levels of active TGFβ1 in bone tissue and bone volume. For every pg/mm^3^ increase in active TGFβ1, bone volume increased by 0.04% (*p*=0.03), when adjusted for OARSI grade, age, sex, and region.

It was also noted that with an increase in one unit of the OARSI grade, osteoclast density decreased by 0.23 cells/mm^3^ (*p*=0.01). However, osteoclast density was significantly higher in CA- regions compared to CA+ regions for 0.9/mm^2^ (*p*=0.02) when adjusted for age and sex. Interestingly, a significant positive association between OARSI grade and volume of osteocyte lacunae, lacunar orientation and shape (sphericity) were identified. For every unit increase in OARSI grade, lacunar volume decreased by 44 µm^3^ (p=0.004), angle of lacunar orientation increased by 2.7 degrees (p=0.008), and sphericity decreased by 0.02 (p=0.04). When adjusted for region, we found that CA- regions had greater lacunar volume by 138.8 µm^3^ and that the angle of lacunar orientation was lower by 8 degrees (p=0.01) compared to CA- regions.

The relationship between OARSI Grade and TGFβ1 levels approached statistical significance (*p*=0.08). Interestingly, it was also noted that TGFβ1 levels also tended to be higher in females compared to males (p=0.08) but this relationship did not reach statistical significance when adjusted for OARSI Grade, Region, and Age. These data are summarised in Table 1.

**Table 1.** Association between active TGFβ1, OARSI score and parameters describing structural, cellular, and molecular tissue characteristics adjusting for sex, age and bone region

| Outcome | Active TGF-B |  | OARSI |  | Region (CA-) |  | Sex (Female) |  | Age |  |
| --- | --- | --- | --- | --- | --- | --- | --- | --- | --- | --- |
|  | P Value | Estimate | P Value | Estimate | P Value | Estimate | P Value | Estimate | P Value | Estimate |
| Active TGF-B | NA | NA | 0.08 | 10.9 | 0.6 | 8.2 | 0.08 | 30.5 | 0.7 | 0.3 |
| TRAP+ Osteoclasts | 0.8 | 0.0001 | **0.01** | **-0.23** | **0.002** | **0.91** | 0.9 | -0.01 | 0.5 | 0.007 |
| TRAP+ Osteocyte | 0.7 | -0.0001 | 0.1 | 0.07 | 0.4 | -0.1 | 0.8 | -0.01 | 0.1 | -0.008 |
| Total_TRAP+ _cells | 0.7 | -0.0001 | 0.1 | -0.15 | **0.008** | **0.81** | 0.8 | -0.03 | 0.8 | -0.001 |
| mRNA_TGF_B1 | 0.7 | -0.0009 | 0.9 | 0.7 | 0.7 | 0.38 | 0.9 | 0.1 | 0.7 | 0.02 |
| mRNA_RANKL | 0.2 | 0.0026 | 0.3 | -0.31 | 0.3 | 0.72 | 0.2 | -0.9 | 0.4 | 0.03 |
| mRNA_OPG | 0.6 | -0.0014 | 0.6 | 0.18 | 0.5 | 0.7 | 0.2 | 1.02 | 0.4 | 0.05 |
| mRNA_TRAP | 0.4 | 0.001 | 0.7 | 0.07 | 0.8 | 0.12 | 0.6 | -0.4 | 0.5 | -0.02 |
| mRNA_OCN | 0.08 | 0.001 | **0.007** | **0.41** | **0.02** | **-0.96** | 0.4 | 0.4 | 0.6 | -0.01 |
| mRNA_COL1 | 0.07 | -0.005 | 0.6 | 0.2 | 0.6 | -0.47 | 0.9 | 0.03 | 0.6 | -0.03 |
| mRNA_ALP | **0.008** | **0.005** | 0.1 | -0.33 | 0.1 | 1.13 | 0.2 | 1.1 | 0.8 | 0.01 |
| mRNA_SMAD3 | 0.3 | -0.002 | 0.3 | 0.37 | 0.3 | -0.92 | 0.8 | -0.2 | 0.5 | 0.03 |
| mRNA_DMP1 | 0.2 | -0.003 | 0.8 | 0.66 | 0.9 | -0.11 | 0.7 | -0.3 | 0.6 | 0.03 |
| BV_TV | **0.03** | **0.04** | 0.09 | 2.5 | 0.2 | 0.09 | 0.3 | -4.8 | 0.5 | 0.1 |
| Tr.Th. | 0.5 | 0.00004 | 0.2 | 0.05 | 0.6 | 0.008 | 0.7 | -0.001 | 0.2 | 0.0002 |
| Tr.No | 0.1 | 0.007 | 0.8 | 0.07 | 0.4 | 0.8 | 0.8 | -0.1 | 0.5 | -0.01 |
| Tr_sep | 0.4 | -0.002 | 0.3 | -0.02 | 0.6 | 0.03 | 0.4 | 0.03 | 0.2 | 0.002 |
| BMD | 0.3 | 2.1 | 0.7 | -53.4 | 0.3 | 484.7 | 0.6 | -228.3 | 0.9 | 1.7 |
| Lacunar_density | 0.3 | 45.47 | 0.1 | 43.4 | 0.4 | 67.2 | 0.8 | 7.9 | 0.9 | -26.1 |
| Lac.Vol | 0.1 | 0.35 | **0.004** | **-44.1** | **0.006** | **138.8** | 0.1 | 32.7 | 0.6 | -0.44 |
| Lacunar Orientation | 0.2 | -0.017 | **0.008** | **2.7** | **0.01** | **-8** | 0.2 | -1.8 | 0.7 | 0.01 |
| Lacunar_SMI | 0.3 | 0.0008 | 0.09 | -0.06 | 0.4 | 0.09 | 0.6 | 0.03 | 0.6 | 0.001 |
| Sauter_diameter | 0.4 | 0.1 | **0.05** | **-79.7** | 0.1 | 176.9 | 0.4 | 163.8 | 0.7 | -348.2 |
| Sphericity | 0.2 | 0.0002 | **0.04** | **-0.02** | 0.2 | 0.04 | 0.6 | 0.1 | 0.7 | 0.0004 |
| Vas_Channels_Dn | 0.8 | 0.029 | 0.7 | 4.5 | 0.4 | 2.3 | 0.7 | -4.3 | 0.4 | -0.5 |
| Vass_diameter | 0.7 | -7.11 | 0.5 | 0.008 | 0.4 | -0.03 | 0.4 | -0.1 | 0.1 | -0.01 |
| Osteocyte cell den. | 0.09 | 0.24 | 0.2 | 21.9 | 0.5 | 34.8 | 0.5 | 24.4 | 0.3 | 2.5 |
| Empty_lacunar_dn. | 0.1 | 0.21 | 0.3 | 16.2 | 0.7 | -17.3 | 0.9 | 49.4 | 0.1 | 3.9 |
| Ratio empty/total_lacunae | 0.1 | 0.26 | 0.5 | 12.4 | 0.6 | 24.7 | 0.4 | 35.9 | 0.1 | 3.7 |
| Arteriolar_density | 0.9 | -0.004 | 0.2 | -40.7 | 0.4 | 90.9 | 0.8 | 15.8 | 0.5 | 2.9 |

## Discussion

The findings of the present study demonstrate that subchondral trabecular bone below severely degraded cartilage (CA-) in human knee osteoarthritis exhibits increased mean concentrations of active TGFβ1, together with an increased ratio of *RANKL*/*OPG* mRNA and increased osteoclast density. Furthermore, the SCB in CA- regions is more sclerotic, hyper-mineralised but with high heterogeneity of the mineral distribution and disorganised collagen matrix with increased osteocyte density and increased vascular canal density, compared to regions of the subchondral trabecular bone below moderately intact cartilage (CA+). These observations are all consistent with increased subchondral bone remodelling being a feature of increased OA disease progression and further suggests that active TGFβ1 could therefore either be a result of these changes or could be playing a driving role. Increased expression of TGFβ1 in human OA bone [34], and evidence for dysregulated expression of TGFβ1 in hip OA [35], have been reported previously. However, the region-specific differences in active TGFβ1 concentrations and their spatial linkage to cartilage and subchondral bone changes in human knee OA are novel, although consistent with the observations made more recently in rodent and guinea pig models of OA [17, 36]. In these animal models, increased activity of TGFβ1 was demonstrated as causal of OA disease, since delivery of anti-TGFβ1 treatments was protective [16, 18].

Our linking here of active TGFβ1 levels to the severity of knee OA degeneration in humans implicates TGFβ1 as a disease target in OA. However, there are several points to make while attempting to compare the human and animal data. While it is tempting to extrapolate from the present study to identify TGFβ1 as a causal agent also in human OA, our observations relate to end-stage disease and therefore cannot inform regarding disease development. We have necessarily compared areas of severe cartilage degeneration with less severe adjacent areas in the same tibial plateau, and we suggest that even the less affected region is unlikely to reflect healthy osteochondral tissue. Despite this potential limitation, it is nevertheless interesting that we see associations between active TGFβ1 levels and structural indices of OA disease. This is particularly so as Zhen *et al*. suggested that increased TGFβ1 activity could be seen early in the mouse anterior cruciate ligament transection (ACLT) OA model, with the structural changes developing later [18]. One interpretation of the mouse data, therefore, might be that anti-TGFβ1 therapy would be ineffective in humans because typically patients do not present with early OA. Although the current data give hope that this might not be the case, it is important to state that there was significant overlap between CA+ and CA- regions in terms of active TGFβ1. This suggests that TGFβ1 might be important in some but not all cases of human knee OA and/or one of a number of molecular contributors to disease progression.

The finding of increased concentrations of active TGFβ1 at sites of severe degeneration in human knee OA raises the question of the significance of TGFβ1 in the subchondral bone at these sites. TGFβ1 has a critical role in bone homeostasis by regulating osteoclast and osteoblast activity [37]. Tight temporal and spatial regulation of the activation of sequestered TGFβ1 is likely essential in healthy bone. Thus, chronic and inappropriately high levels of TGFβ1 activation is likely deleterious in bone [14], as it is in other tissues [38]. *In vitro* experiments show that an excess of TGFβ1 alters the ratio of type I collagen alpha 1 and alpha 2 chains, leading in turn to under-mineralisation of osteoblast cell layers, since neutralisation of TGFβ1 corrects these abnormalities [39]. Chronic overloading of the knee joint is a known major contributor to the development of OA [40]. Rys *et al*. have compiled evidence for a model, in which TGFβ1 regulates the cellular response to loading in the skeleton, in both cartilage and bone compartments [41]. According to this model, overloading results in both excess TGFβ1 and dysregulated responses to it. As reviewed by Crane and Cao [37], increased levels of active TGFβ1 can result from inappropriate bone loading, increased osteoclastic resorption, mutations in the TGFβ1 molecule itself or in extracellular matrix molecules that normally bind TGFβ1, and for reasons not yet understood, and these can all lead to dysregulated recruitment of osteoblast progenitors and abnormal bone remodelling.

In the present study, we indeed found an association between increased concentrations of activated TGFβ1 protein and osteoclast density and activity. We also found TRAP^+^ osteocytes present in both CA+ and CA- subchondral bone, a phenomenon which has been reported previously in association with perilacunocanalicular remodelling [42]. Demineralisation surrounding osteocytes in CA- regions suggests that osteocytes contribute to bone matrix remodelling in knee OA. Factors known to be involved in this process include MMP13 [43], sclerostin, carbonic anhydrase II and cathepsin K [42]. It was recently reported that, in both mouse and human subchondral bone, lower expression of MMP13 and cathepsin K due to osteocyte dysfunction is site-dependent and associates closely with suppression of perilacunar remodelling. The result is collagen fibril disorganisation and hyper-mineralisation of the subchondral bone matrix [31] [22]. In a separate study, it was demonstrated that TGFβ1 signalling has an intrinsic role in osteocyte function [12]. These results are consistent with our finding of region-dependent collagen disorganisation and hyper-mineralisation of subchondral bone matrix, which correlate with increased levels of active TGFβ1.

It is of interest that the ratio of *RANKL*/*OPG* mRNA was the only molecular index to show an association with the active TGFβ1 level in the subchondral bone samples. This is likely to be a function of the expression of *RANKL* mRNA species in osteocytes, since (a) osteocytes are the most abundant cells in bone and therefore most likely the major contributors to the mRNA assayed in this study, and (b) osteocytes are reported to be the major producers of *RANKL* in adult bone [44]. Evidence also supports osteocytes as the most important mechano-responsive cells in bone [45], with the greatest potential to link mechanical overloading in OA to a cellular response. Interesting also, is a report that mechanosensation of subchondral bone require osteocytic TGFβ1 signalling [33] and that compressive stress changed the effect of TGFβ1 on RANKL and OPG expression in cement-oblasts and osteoblasts [46].

Consistent with our own data and the findings of others [5, 47], we found increased osteocyte density and vascular density in the SCB immediately below areas of severe cartilage degradation (CA-). These observations align with those from the mouse ACLT study already cited [18]. It was shown in those studies that increased TGFβ1 activity was causal in the development of both abnormal bone formation and increased angiogenesis. [17, 18]. It has also been reported that TGFβ1 directly promotes angiogenesis in endothelial progenitor cells [13, 48]. Again, our human knee OA findings are consistent with the mouse data and support a hypothesis that abnormal remodelling and increased angiogenesis in human OA might be due to increased TGFβ1 activity. We propose that the increased osteocyte density and a decrease in empty osteocyte lacunae is related to increased bone remodelling rate at sites of more severe OA.

Another intriguing possibility is the relationship between active TGFβ1 and pain associated with knee OA in the SCB [6]. TGFβ1 has been shown to stimulate the pain mediator nerve growth factor (NGF) by synovial cells and chondrocytes in knee OA [49]. A study from our group showed that osteocytes derived from the knee SCB in patients undergoing total knee replacement surgery, could also produce NGF, implicating them in the pain response [50]. It remains to be seen if the regional difference in TGFβ1 levels is connected to the expression of neurotropic factors including NGF and connected to pain.

A strength of the current study is the use of human OA osteochondral samples to investigate the structural, cellular and molecular mechanisms of OA progression, rather than the use of animal models or cultured cells. However, as already noted, this presents limitations, including the inability to obtain a healthy non-OA sample cohort for comparison and the establishment of causality.

In conclusion, this study supports the hypothesis that an increased level of active TGFβ1 occurs in human knee OA and associates spatially with disease severity. The increase in TGFβ1 activity was associated, at the tissue and cellular level, with elevated markers of bone remodelling and the ratio of *RANKL*/*OPG* mRNA, and vascular changes. The study therefore suggests that TGFβ1 could be a therapeutic target in at least a subset of human knee OA patients, to prevent or reduce disease progression.

## Acknowledgements

The authors acknowledge the Paul Scherrer Institut, Villigen, Switzerland for provision of the synchrotron radiation beamtime at the TOMCAT beamline of the Swiss Light Source and would like to particularly thank to Dr. Elena Borisova and Dr Christian Schlepuetz for their support during and after beamtime at TOMCAT.

We also acknowledge travel funding provided by the International Synchrotron Access Program (ISAP) managed by the Australian Synchrotron, part of ANSTO, and funded by the Australian Government.

The authors wish to thank Dr Stuart Callary and Dr Roumen Stamenkov for assistance in obtaining the tibial plateau specimens, Associate Professor Egon Perilli and Dr Jasvir Bahl for insightful advice how to obtain and visualise data from Synchrotron micro-CT images. In addition, we are grateful to Dr Catherine Stapledon for assistance in developing the method for reverse transcription polymerase chain reaction (RT-PCR), Ms Yea Rin Lee and Ms Halimatul Masrini Binti Haji Masri for her help preparing sections for histomorphometry analysis and histological staining and Ms Halah Al-Dulaimi for her help with histological staining.

The authors wish to acknowledge support from Adelaide Microscopy at The University of Adelaide, especially Dr Agatha Labrinidis, for assisting with developing the method to obtain data from Synchrotron micro CT images and Dr Jane Sibbons for her support during polarised light microscopy

## Author contributions

All authors meet the criteria for authorship. DM designed the study, performed all experiments and analysis of the results, interpreted the data, and wrote the manuscript. RQ performed statistical analysis and developed MATLAB code/programs for data collection, interpreted the data and approved the manuscript, DF and JK, XC, BS, GA designed the study, interpreted the data and provided overall supervision. All authors read and approved the manuscript.

