## Supplementary figures and images for "Elevated levels of active Transforming Growth Factor β1 in the subchondral bone relate spatially to cartilage loss and impaired bone quality in human knee osteoarthritis"

### Suplementary Figure 1

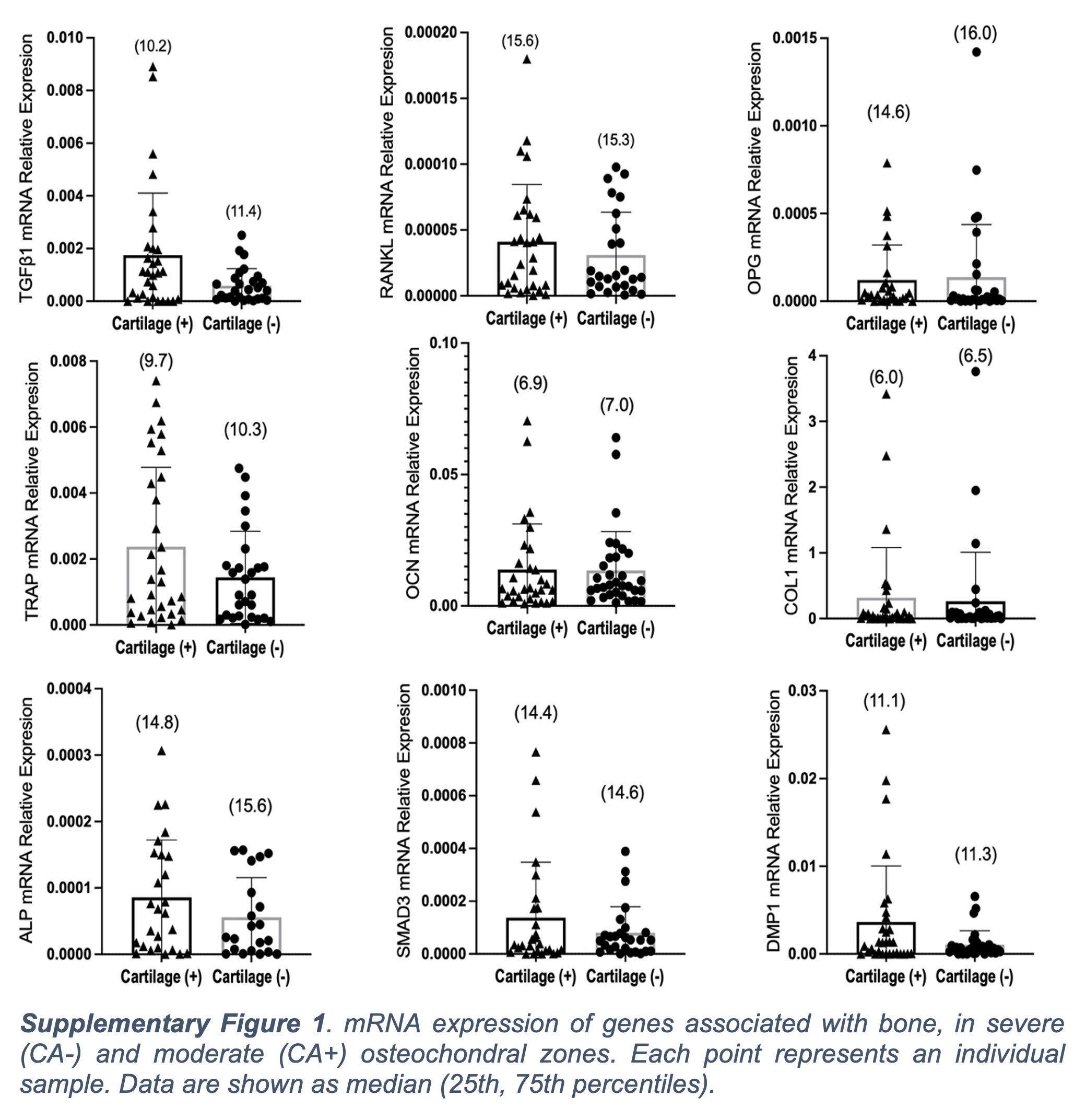
