## Supplementary material for "Elevated levels of active Transforming Growth Factor β1 in the subchondral bone relate spatially to cartilage loss and impaired bone quality in human knee osteoarthritis": Suplementary Method

### 1. Protein assay

Core biopsies used to determine the concentration of TGFβ1 were cleaned from bone marrow by several wash in sterile PBS prior crushing in 2 ml of PEB (2 mM EDTA, 0.2% BSA in PBS pH 7.4) using a mortar and pestle. Further, liberated cells were separated from the bone debris by filtering through a 70µm filter and the cell suspension was centrifuged at 13000xg for 15 min. The clarified supernatant was divided into two sample tubes to quantify active TGFβ1 and total (active + latent) TGFβ1 using the Quantikine Human TGFβ1 Immunoassay (DB100B, R&D Systems, Minneapolis, MN, USA), according to the manufacturer’s instructions. Specifically, to determine the total (active + latent) TGFβ1 levels, the samples were acidified to pH 2.0 using 1N HCl and then neutralised by adding 20 μl of 1.2 N NaOH/0.5 M HEPES (pH 7.2-7.6) before addition to the assay plate. The optical density was determined within 30 minutes using a MR7000 microplate reader (Dynatech Laboratories, Guernsey, Channel Islands) set to 450 nm.

### 2. Reverse transcription polymerase chain reaction (RT-PCR)

Bone core biopsies were cleaned of bone marrow by several washes in sterile PBS and then finely crushed in liquid nitrogen, using a sterile mortar and pestle. The crushed bone was transferred to sterile 1.5 ml centrifuge tubes, and 1 ml of cold Trizol reagent (Life Technologies, Gaithersburg, MD) was added. The samples were frozen at -80°C overnight, thawed, then total RNA prepared as per the manufacturer’s instructions. The yield and purity of RNA were tested using a Nanodrop microvolume spectrophotometer (Thermo Fisher Scientific, Delaware, USA). RNA was reverse transcribed using Superscript TM IV (Thermo Fisher Scientific). Gene expression was analysed by real-time RT-PCR using SYBR Green Fluor qPCR Mastermix (Qiagen, Limburg, The Netherlands) in a CFX Connect Real Time PCR System (Bio-Rad, Hercules, CA, USA). RT-PCR was performed for transforming growth factor beta 1 (*TGFB1*), small mothers against decapentaplegic 3 (*SMAD3*,) tartrate-resistant acid phosphatase (*TRAP*/*ACP5*), receptor activator of NF-κB ligand (*RANKL*/*TNFSF11*), osteoprotegerin (*OPG*/*TNFRSF11B*), dentin matrix acidic phosphoprotein 1 (*DMP1*), tissue non-specific alkaline phosphatase (*TNAP*), osteocalcin (*OCN*) and collagen type I alpha 1 (*COL1A1*) genes, and mRNA levels were expressed relative to mRNA encoding the housekeeping gene *18S*. Oligonucleotide primer sequences for each of these PCR products are shown in Table 1. The relative quantification method was used to quantify the mRNA levels ^56^.

**Table 1** List of genes and primer sequences

| Gene | Primer sequence (5´ – 3´) |
| --- | --- |
| *18S* | F – GCG TTG ATT AAG TCC CTG CC |
|  | R – CAC CTA AGG AAA CCT TGT TAC GAC |
| *TGFB1* | F – GAC ACC AAC TAT TGC TTC AG |
|  | R – AGA AGT TGG CAT GGT AGC CC |
| *SMAD3* | F – TTC AAC AAC CAG GAG TTC GC |
|  | R – TAC TGG TCA CAG TAT GTC TC |
| *TRAP* | F – GTG CAG ACT TCA TCC TGT CTC TA |
|  | R – AAT ACG TCC TCA AAG GTC TCC |
| *RANKL* | F – TCA GCC TTT TGC TCA TCT CAC TAT |
|  | R – CCA CCC CCG ATC ATG GT |
| *OPG* | F – GTC CAC AAG AAC AGA CTT TCC AG |
|  | R – CTG TTT TCA CAG AGG TCA ATA TCT T |
| *OCN* | F – TGA GAG CCC TCA CAC TCC TC |
|  | R – ACC TTT GCT GGA CTC TGC AC |
| *DMP1* | F – GAT CAG CAT CCT GCT CAT GTT |
|  | R – AGC CAA ATG ACC CTT CCA TTC |
| *ALP* | F – TGC TCC CAC GCG CTT GTG CCT GGA |
|  | R – CTG GCA CTA AGG AGT TAG TAA G |
| *COL1A1* | F – AGG CCT CCA ACG AGA TCG AGA TCC G |
|  | R – TAC AGG AAG CAG ACA GGG CCA ACG TCG |

### 1. Synchrotron radiation micro-computer tomography (SRμCT)

Synchrotron tomographic X-ray experiments were carried out at the X02DA TOMCAT beamline of the Swiss Light Source (SLS) facility at the Paul Scherrer Institute (PSI). Prior to imaging, bone core biopsies of trabecular bone were fixed in formaldehyde for 24h, several times rinsed in 75% ethanol and mechanically embedded in paraffin to be protected from overheating damage during scanning time. Bone cores were scanned with a monochromatic beam, with energy 20 keV used to acquire tomography scans. The X-ray imaging system consisted of a microscope (Optique Peter) with 4x magnification objective and LuAG:Ce 20μm scintillator coupled to a pco.edge 5.5 detector with a pixel size of 1.63 μm and field of view 4.2x3.5 mm^2^. For each sample 1800 projections uniformly distributed over 180 degrees were taken with 120 ms exposure each, resulting in scan times of approximately 6 minutes. Tomographic reconstructions were done on the TOMCAT cluster ^57^ using phase retrieval method from a single defaced image ^58^ and the GridRec algorithm ^59^. Final reconstructed images were saved as 16 bit TIFF image stacks.
